# Neuroprotective Efficacy of the Glucocorticoid Receptor Modulator PT150 in the Rotenone Mouse Model of Parkinson’s Disease

**DOI:** 10.1101/2024.04.12.589261

**Authors:** Amanda S. Latham, Savannah M. Rocha, Casey P. McDermott, Philip Reigan, Richard A. Slayden, Ronald B. Tjalkens

**Affiliations:** Department of Environmental and Radiological Health Sciences, Colorado State University, Fort Collins, CO 80523; Department of Pharmaceutical Sciences, Skaggs School of Pharmacy and Pharmaceutical Sciences, University of Colorado Anschutz Medical Campus, Aurora, CO, United States; Department of Microbiology, Immunology and Pathology, Colorado State University, Fort Collins, CO 80523

## Abstract

Parkinson’s disease (PD) is the most common neurodegenerative movement disorder worldwide. Current treatments for PD largely center around dopamine replacement therapies and fail to prevent the progression of pathology, underscoring the need for neuroprotective interventions. Approaches that target neuroinflammation, which occurs prior to dopaminergic neuron (DAn) loss in the substantia nigra (SN), represent a promising therapeutic strategy. The glucocorticoid receptor (GR) has been implicated in the neuropathology of PD and modulates numerous neuroinflammatory signaling pathways in the brain. Therefore, we investigated the neuroprotective effects of the novel GR modulator, PT150, in the rotenone mouse model of PD, postulating that inhibition of glial inflammation would protect DAn and reduce accumulation of neurotoxic misfolded ⍺-synuclein protein. C57Bl/6 mice were exposed to 2.5 mg/kg/day rotenone by intraperitoneal injection for 14 days, immediately followed by oral treatment with 30 mg/kg/day or 100 mg/kg/day PT150 in the 14-day post-lesioning incubation period, during which the majority of DAn loss and α-synuclein (α-syn) accumulation occurs. Our results indicate that treatment with PT150 reduced both loss of DAn and microgliosis in the nigrostriatal pathway. Although morphologic features of astrogliosis were not attenuated, PT150 treatment promoted potentially neuroprotective activity in these cells, including increased phagocytosis of hyperphosphorylated α-syn. Ultimately, PT150 treatment reduced the loss of DAn cell bodies in the SN, but not the striatum, and prohibited intra-neuronal accumulation of α-syn. Together, these data indicate that PT150 effectively reduced SN pathology in the rotenone mouse model of PD.

## Introduction

Parkinson’s Disease (PD) is the second most common neurodegenerative disorder, with an estimated six million individuals that are directly affected worldwide [1, 2]. The impact of PD may be underestimated due to challenges in accurate diagnostic modalities of the disease, making the current and projected disease burden higher than previously reported [3]. PD is characterized clinically by motor symptoms, including resting tremor, rigidity, bradykinesia, and postural instability, in addition to non-motor features [4]. Non-motor symptoms occur early in prodromal disease, preceding motor symptoms by as much as decades. These non-motor disturbances include anxiety, depression, and gastrointestinal dysfunction [4]. The etiology of PD remains largely unknown, and it is primarily considered an age-related disorder, but epidemiological and experimental evidence suggests that risk factors include genetic mutations and exposure to environmental contaminants. Pesticides, such as rotenone, have been associated with increased risk of developing PD [5]. Rotenone is a mitochondrial complex I inhibitor that results in high levels of reactive oxygen species (ROS), culminating in oxidative stress and disrupted cellular signaling [6]. Ultimately, exposure to rotenone causes selective loss of dopaminergic neurons (DAn) in the substantia nigra pars compacta (SNpc) and retraction of terminal axonal projections within the striatum (ST). This, along with chronic neuroinflammation and the hyperphosphorylation and subsequent aggregation of α-synuclein (α-syn) into intra-neuronal Lewy bodies, comprise the primary neuropathological features of PD. We previously demonstrated that in mice exposed to rotenone for 14 days by intraperitoneal injection, neuroinflammatory activation of glia and widespread aggregation of ⍺-synuclein occurred prior to loss of DAn in the SNpc, indicating that these pathologic changes were key drivers of neuronal injury [7].

During neuroinflammation, microglia and astrocytes polarize from resting states to neurotoxic or pro-inflammatory phenotypes in response to environmental, pathogenic, or genetic stressors. These activated states aid in restoring brain homeostasis during stress by recruiting immune cells, removing pathogens and misfolded proteins through phagocytosis, and by secreting neurotrophic factors [8]. Despite these beneficial activities associated with adaptive glial reactivity, prolonged cellular activation creates a pro-inflammatory environment that chronically damages the brain [9]. The dysregulation of neurotransmitters and the production of ROS resulting from glial activation promotes neuronal dysfunction and neurodegeneration. In addition, neuroinflammation is hypothesized to contribute to the propagation of neurotoxic α-syn, as these pathologies are dynamically interconnected, which exacerbates the formation of Lewy bodies and worsens neuropathology [10]. Therefore, modulation of glial activation and subsequent neuroinflammation remains a promising target for therapeutic intervention.Notable inflammatory signaling pathways mediated by glia that have been identified in PD include nuclear factor-kappa B (NF-κB) and nucleotide-binding oligomerization domain, leucine-rich repeat-containing protein 3 (NLRP3). Activation of these pathways increases production of pro-inflammatory cytokines [11]. One potential avenue for controlling this pathological response is through the glucocorticoid receptor (GR), which is found ubiquitously in all parenchymal cells of the brain, particularly in astrocytes and microglia. GR has the capacity to regulate inflammatory signaling, dependent on the brain region, cell type, and the physiological context involved. Upon activation of the cytosolic GR by glucocorticoid (GC) hormones, it releases from chaperone proteins and translocates to the nucleus. There, it binds glucocorticoid response elements (GREs), thereby altering the transcription of inflammatory genes. The GR-GC complex can also interact with cytosolic proteins for post-translational regulation of inflammation. High quantities of GRs on both DAn and glia in the midbrain have been identified in rodent models, which have the capacity to trigger functional changes in the dopamine system [12]. GCs also have an expansive history of clinical use as anti-inflammatories for numerous chronic inflammatory diseases, such as multiple sclerosis and rheumatoid arthritis [13]. Altogether, this suggests that targeting the GR is a potential therapeutic avenue for treating the neuroinflammation associated with PD.

In addition to its anti-inflammatory potential, GR has been directly linked to the neuropathology associated with both the aged brain and neurodegenerative diseases, including PD. GR and the GC hormone, cortisol in humans or corticosterone in rodents, are altered during aging and disease. Not only is GR function selectively reduced with age, murine models of PD exhibit fewer GRs in the SN [14, 15]. Similarly, analysis of post-mortem PD midbrain tissue demonstrates a significant decrease in the number of astrocytes expressing nuclear GR, which diminishes its anti-inflammatory capacity [16]. Moreover, an imbalance of the hypothalamic-pituitary-adrenal axis described in PD patients and in rodent models results in chronically increased cortisol/corticosterone levels [17–19]. High cortisol is linked to cognition decline and risk for neurodegenerative disease, as well as neuroinflammatory effects. It is unclear whether this is simply correlation or a causal effect, although studies show that stress, which induces the metabolism of cortisol, can worsen the motor symptoms associated with PD [20]. GRs can become inactivated or desensitized in the presence of chronically increased cortisol/corticosteroid, therefore disrupting the ability of GR to regulate inflammation [17, 18].

The function of GR in the central nervous system (CNS) is complex, as its actions cannot be uniformly classified as anti-inflammatory, but targeting GR has shown neuroprotective effects. PD models show that inactivating astrocytic GRs exacerbates DAn loss in the substantia nigra (SN) compared to controls and increases both glial reactivity and levels of pro-inflammatory cytokines [15, 21]. In experimental models of PD, activating microglial GRs impedes neurotoxic glial activation, reducing DAn degeneration [18]. Limiting neuroinflammation in microglia-specific NF-κB knockout mice reduced reactive gliosis and preserved the number of DA neurons in the SNpc [22]. However, complete inhibition of glial activation can exacerbate damage in response to stress by preventing secretion of trophic factors, limiting repair mechanisms, and preventing removal of misfolded proteins, which are necessary responses to stress or injury [23]. Therefore, therapeutics that modulate the neuroinflammatory signaling from glia, but do not completely attenuate their activity, may be more protective.

In this study, we examined the therapeutic efficacy of the GR modulator, (11b,17b)-11-(1,3-benzodioxol-5-yl)-17-hydroxy-17-(1-propynyl)-estra-4,9-dien-3-one (PT150), in preventing the neuropathology associated with rotenone neurotoxicity [24]. We previously reported on the anti-inflammatory effects of PT150 during viral infection, as well as its capacity to modulate GR-dependent gene expression [25]. Based on these data, we hypothesized that PT150 would effectively decrease glial inflammation, prohibit the aggregation of α-syn, and reduce the degeneration of DAn in the SNpc. Following exposure to rotenone (2.5mg/kg/day for 14 days), daily oral administration of PT150 for two weeks reduced glial reactivity and preserved dopaminergic soma in the SN, although no protective effects were seen in axonal projections in the ST. Additionally, PT150 treatment decreased intraneuronal α-syn accumulation within DAn cell bodies in the SNpc. These data suggest that pharmacologic modulation of GR is neuroprotective in the rotenone model of PD by altering pro-inflammatory glial responses.

## Materials and Methods

### Molecular Modeling

PT150 was docked into the co-activator site and the steroid binding site of the crystal structures of the ligand-binding domain of the glucocorticoid receptor (PDB: 3CLD and PDB1NHZ) [26, 27] using the Glide module within Schrödinger (Release 2020-2, Schrödinger LLC, New York, NY) [28–30]. Prior to docking, the water molecules were removed, and the proteins were prepared by assigning bond orders, adding hydrogens, and repairing any side chains or missing amino acid sequences. To complete protein preparation, a restrained minimization of the protein structure was performed using the default constraint of 0.30Å RMSD and the OPLS_2005 force field [31]. The prepared proteins were subjected to SiteMap analysis [30], that identified the available binding sites in the ligand binding domains of the glucocorticoid receptors. Docking grids were generated using Receptor Grid Generation. PT150 was prepared using LigPrep by generating possible states at the target pH 7.0 using Epik and minimized by applying the OPLS_2005 force field [31]. Molecular docking simulations were performed by targeting each potential binding site for PT150 using the Glide ligand docking module in XP (extra precision) mode and included post-docking minimization [29].

### Animals Procedures and Sample Collection

Experiments consisted of male and female C57Bl/6J mice three months of age (*N* = 8/group) randomly assigned to experimental groups. Animals were housed at the Colorado State University Laboratory Animal Resources facility accredited by the American Association for Accreditation of Laboratory Animal Care (AAALAC). All animal experiments were performed in accordance with the National Research Council’s Guide for the Care and Use of Laboratory Animals and were approved by the Institutional Animal Care and Usage Committee (IACUC) at Colorado State University. Experimental animals were housed under constant temperature and humidity conditions (21° ± 2 °C temperature and 30 ± 5 % humidity) and a 12-hour light/12-hour dark cycle was used. Mice had *ad libitum* access to standard pelleted food and water and were monitored using a clinical scoring system for signs of morbidity.

Rotenone was diluted to a final working solution daily in medium chain-triglyceride, miglyol + 2% dimethyl sulfoxide (DMSO) [32]; a 50X stock solution was prepared in 100% DMSO every 48 hours and stored at −20°C. The head space of the vial was purged with nitrogen to prevent oxidation of the compound. Working concentrations were prepared by diluting 50X stock solutions in Miglyol 812 as previously described [7, 22]. Rotenone was delivered by intraperitoneal (i.p.) injection (2.5 mg/kg/day), as previously determined [7]. Vehicle groups received an equivalent volume of 100% miglyol by i.p. injection. Mice were weighed daily and a dosage of 2 μL/g body weight rotenone or vehicle was measured using a 50 μL Hamilton syringe, which was then transferred to an insulin syringe for i.p. injection once daily for 14 days. Hamilton syringes were cleaned daily in 10% bleach for 10 minutes, followed by 70% ethanol, then sterile water to prevent precipitate buildup within the needle.

Animals were randomly assigned to the following experimental groupings (*N* = 8 animals/group): vehicle (Miglyol), 2.5 mg/kg rotenone + vehicle (Miglyol), 2.5 mg/kg rotenone + 30 mg/kg PT150, and 2.5 mg/kg rotenone + 100 mg/kg PT150. Treatment dosage of the experimental drug, PT150 (supplied by Palisadse Therapeutics/Pop Test Oncology, LLC), was calculated using dosing schemes and toxicity data from human clinical trials, which treated with 500 mg of PT150, or approximately 7 mg/kg. Calculations normalizing to body surface area determined that a dose of approximately 86 mg/kg in mice equates to a Human Equivalent Dose (HED) of 7 mg/kg [24, 33]. Thus, two doses of PT150 were evaluated for the duration of the study, a low dose of 30 mg/kg and high dose of 100 mg/kg. PT150 was dissolved in miglyol and delivered by oral gavage at 8μL/g body weight once daily for 14 days after the conclusion of rotenone dosing. Vehicle groups received 100% miglyol by oral gavage. At the conclusion of the study, animals were euthanized by decapitation under isoflurane anesthesia and tissues were collected for histopathology by fixation in 10% neutral buffered formalin.

### Histopathological Processing and Immunofluorescent Staining

Brains isolated from mice were fixed for 72 hours in 10% neutral buffered formalin before being transferred to Colorado State University’s Veterinary Diagnostic Laboratory for tissue sectioning. Paraffin-embedded brain tissue was sectioned at 5µm thickness and mounted onto charged slides. Brain sections were deparaffinized and stained for immunofluorescence detection usingthe Leica Bond RX_m_ automated robotic staining system (Leica Biosystems, Nussloch GmbH). Antigen retrieval was performed using Bond Epitope Retrieval Solution 1 for 20 minutes at 60°C. Sections were permeabilized (0.1% Triton X in 1X tris-buffered saline/TBS) and blocked in 1% donkey serum or 1% donkey + goat serum in 1X TBS. Primary antibodies were diluted in 1X TBS and incubated for 1 hour per antibody. The following antibodies were used: rabbit anti-tyrosine hydroxylase (TH) (1:500; Millipore, cat #: AB152), mouse anti-neuronal nuclei [34] (1:200; Abcam, Cat #: ab279296), mouse anti-glial fibrillary acidic protein (GFAP) (1:1000; Abcam, Cat #: ab4648), rabbit anti-S100 Calcium Binding Protein B (S100β) (1:750; Abcam, Cat #: ab41548), rat anti-complement component 3 (C3) (1:250; Abcam, Cat #: ab11862), goat anti-ionized calcium binding adaptor molecule 1 (Iba-1) (1:50; Abcam, Cat #: ab5076), mouse anti-α-syn phosphorylation at serine position 129 (p129) (1:100; FUJIFILM Wako Chemicals, Cat #: 015-25191), and rabbit anti-α-syn (1:100; Abclonal, Cat #: A7215). Sections were stained with Hoescht 33342 (DAPI) (diluted 1:5000 in PBS; Invitrogen, Cat #: H3570) and mounted on glass coverslips in ProLong Gold Antifade hard set mounting medium (Fisher Scientific, Cat #: P36930). Slides were kept at room temperature for 24 – 48 hours to allow mounting medium to harden, and then stored at 4°C prior to imaging.

### Immunofluorescence Imaging and Protein Quantification

Whole-tissue immunofluorescence montage images were captured using an Olympus VS200 slide scanning fluorescent microscope equipped with a Hamamatsu ORCA-fusion CMOS digital camera and collected using Olympus CellSens software. Full slide images were acquired using an Olympus X-line Apochromat 20X air objective (0.8 N.A.). All images were obtained using the same exposure time, light source intensity, camera gain, and filter application per channel. Brain regions were identified anatomically and regions of interest (ROIs) were manually drawn using Olympus CellSens software. The manual or adaptive thresholding features of the Count and Measure function of Olympus CellSens software were used to quantify immunofluorescence images for S100β^+^ cell number and GFAP^+^ area. Object filtering was used to remove non-cellular objects from the analysis. Intracellular protein expression was analyzed by using manual thresholding on the Count and Measure function of Olympus CellSens software to identify cells (S100β^+^ or GFAP^+)^, creating an ROI from each detected object, and then analyzing the fluorescence intensity of each individual cellular ROI.

### Stereological and Immunofluorescent Quantification of Dopaminergic Neurons in the SN and ST

Quantification of DAn was conducted using unbiased stereological methods adapted from our previous reports [7, 22, 35, 36]. Six sections per animal spanning the entire substantia nigra (SN) were analyzed blindly by a single investigator. Regions of interest (ROIs) were drawn manually using TH^+^ immunolabelling in combination with the Allen Brain Atlas (Allen Institute for Brain Science, Seattle, WA, USA) to identify the SN and surrounding anatomical landmarks. TH^+^ and NeuN^+^ cells were quantified by using adaptive thresholding in the Count and Measure feature on the Olympus CellSens software, followed by object filtering. Quantitative stereological analysis using the motorized stage method was performed as previously described[37]. Striatal sections were immunostained for TH and fluorescence intensity was analyzed by manually drawing an ROI and using manual thresholding on the Count and Measure function of Olympus CellSens software as previously reported [7].

### Morphological Characterization of Glia

Morphometric analysis was performed using Imaris image analysis software (version 9.8.2, Bitplane Imaris, South Windsor, CT, USA). Four randomized 400 × magnification images spanning the entirety of the substantia nigra were taken using an Olympus X-line Apochromat 40X air objective (0.95 N.A.). The Filament Tracing module was used to identify GFAP^+^ astroglial and Iba-1^+^ microglial processes. A total sum of processes per cell (filament length [sum]), branch number per cell (filament number of dendrite terminal points), and overall volume of processes per cell (filament volume [sum]), were utilized to determine morphometric changes present within each animal. Skeletonized renderings of glial cells were performed by using Imaris software (version 9.8.2, Bitplane Imaris, South Windsor, CT, USA) using high-magnification images.

### Statistical Analysis

All data is shown as mean +/− SEM, unless otherwise noted. Experimental values were analyzed using a ROUT (α=0.05) test, and significant outliers there removed from the data set. Differences between each experimental group were analyzed using a one-way ANOVA with Tukey’s post hoc test. All statistical analysis was conducted using Prism. Significance is denoted as * *p* < 0.05, ** *p* < 0.01, *** *p* < 0.001, **** *p* < 0.0001.

## Results

### PT150 is likely an allosteric modulator of the glucocorticoid receptor through interactions with the co-activator binding domain

Through computational-based molecular modeling, binding of PT150 to the co-activator and steroid binding sites of the ligand-binding domain of the glucocorticoid receptor was simulated using two separate protein structures (PDB: 3CLD and PDB: 1NHZ). First, PT150 was docked against the co-activator site and steroid binding site of the PDB: 3CLD crystal structure using the Glide module within Schrödinger (Release 2020-2, Schrödinger LLC, New York, NY). The 2.84Å PDB: 3CLD glucocorticoid receptor crystal structure has an intact ligand-binding domain encompassing 259 residues and co-crystallized with fluticasone in the steroid binding site (fluticasone was removed prior to docking). A docked pose for PT150 was obtained in the co-activator site where the entire dodecahydrocyclopentaphenanthren-3-one steroid-like nucleus occupies the hydrophobic site with the benzodioxole ring orientated out of the pocket (**Figure 1A** and **B**). No anchoring H-bond interactions were observed between PT150 and the co-activator site (**Figure 1B**). Interestingly, no docked pose for PT150 was identified in the steroid binding site; however, the 2.30Å PDB: 1NHZ glucocorticoid receptor crystal structure was co-crystallized with RU486, a structural analog of PT150, in the steroid binding site. The co-crystal structure of RU486 in the steroid binding site of the ligand-binding domain of the glucocorticoid receptor (PDB: 1NHZ), suggests that the binding of PT150 would also be possible within the steroid binding pocket of the ligand binding domain. However, the helix that forms at the interface of both the steroid site and the co-activator site, defining these sites, is not complete due to a 9 amino acid deletion. The consequence of the deletion of these amino acids is that both the co-activator and steroid binding sites lose their integrity making the steroid site more accessible and results in significant distortions in the conformation of the steroid-binding site and the overall tertiary structure of the protein. Despite this issue in the 1HNZ crystal structure, there is still potential for PT150 to bind the steroid binding site of the glucocorticoid receptor as the dynamic movement of the protein cannot be completely recapitulated in flexible docking; however, our docking studies support that PT150 has the potential to bind to the co-activator site and modulate glucocorticoid receptor-mediated transcriptional activity and that this site may be more accessible than the steroid binding site.

**Figure 1.**
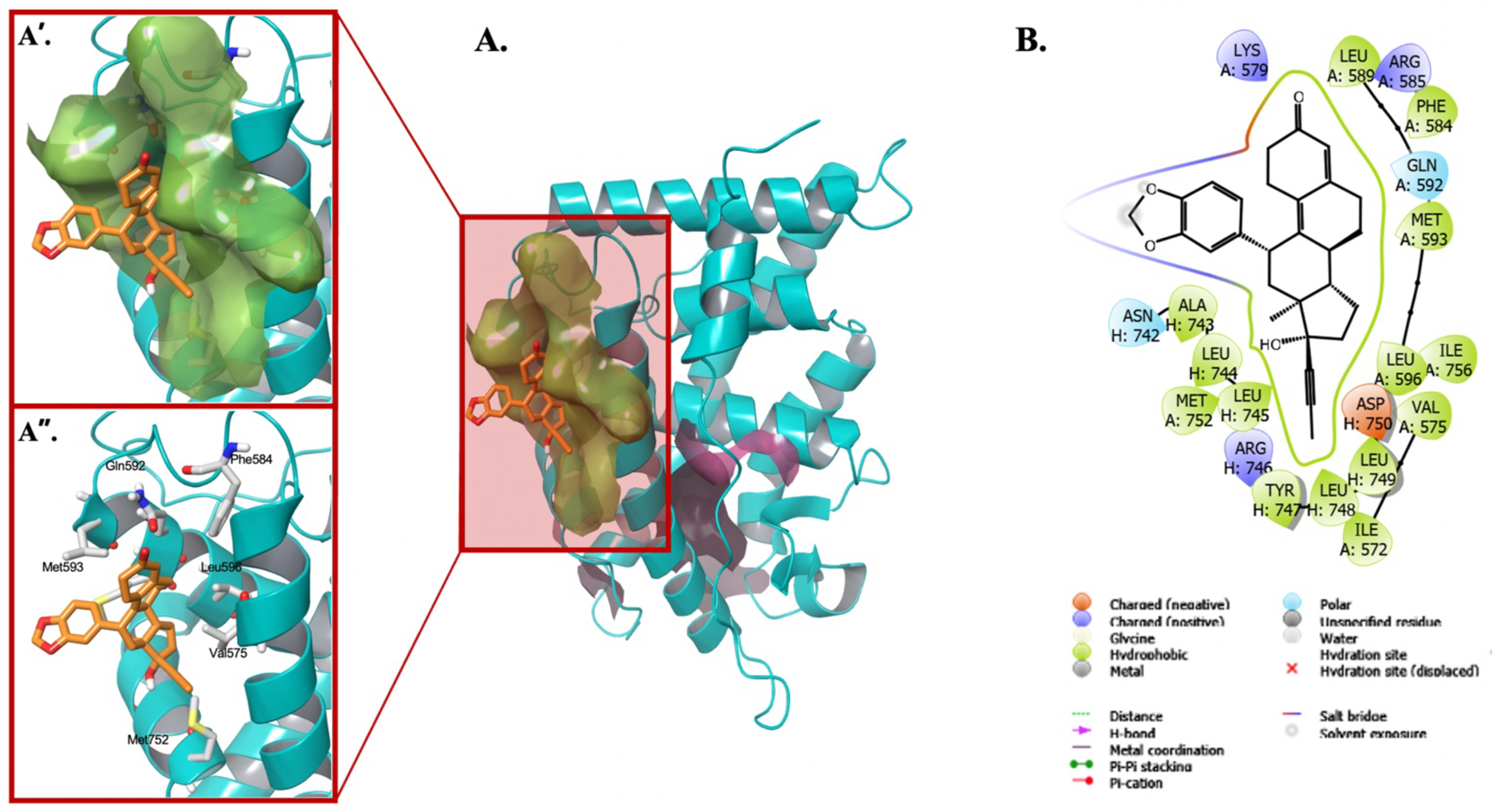
Computational-based molecular docking of PT150 interactions with the ligand binding domain of the human glucocorticoid receptor. PT150 was docked into the co-activator site and the steroid binding site of the crystal structures of the ligand-binding domain of the glucocorticoid receptor. Restrained minimization of the protein structure was performed and prepared proteins were subjected to SiteMap analysis to identify the available binding sites. The interaction between PT150 and the glucocorticoid receptor demonstrates increased affinity for the co-activating domain (A), with the benzodioxole ring orientated out of the pocket (A’, A”). No anchoring H-bond interactions were observed between PT150 and the co-activator site (B).

### Treatment with 30 mg/kg PT150 did not result in unexpected animal mortality

Two doses of PT150 were evaluated for the duration of the study, 30 mg/kg and 100 mg/kg. Adult male and female C57Bl/6 mice were exposed to a daily intraperitoneal (i.p.) dose of 2.5 mg/kg rotenone or vehicle for 14 days, followed by daily treatment with PT150 or vehicle for 14 days by oral gavage (**Figure 2A**). Signs of morbidity or mortality were monitored daily for each animal. Upon conclusion of the study, animals that received rotenone in combination with 100 mg/kg PT150 had a 50% mortality (**Figure 2B**). There was no difference in mortality between the rotenone-exposed animals that received 30 mg/kg PT150 or vehicle by the conclusion of the study (**Figure 2B**). Due to adverse health effects associated with rotenone + 100 mg/kg PT150 treatment, only the rotenone + 30 mg/kg PT150-treated group was analyzed for the remainder of the study.

**Figure 2.**
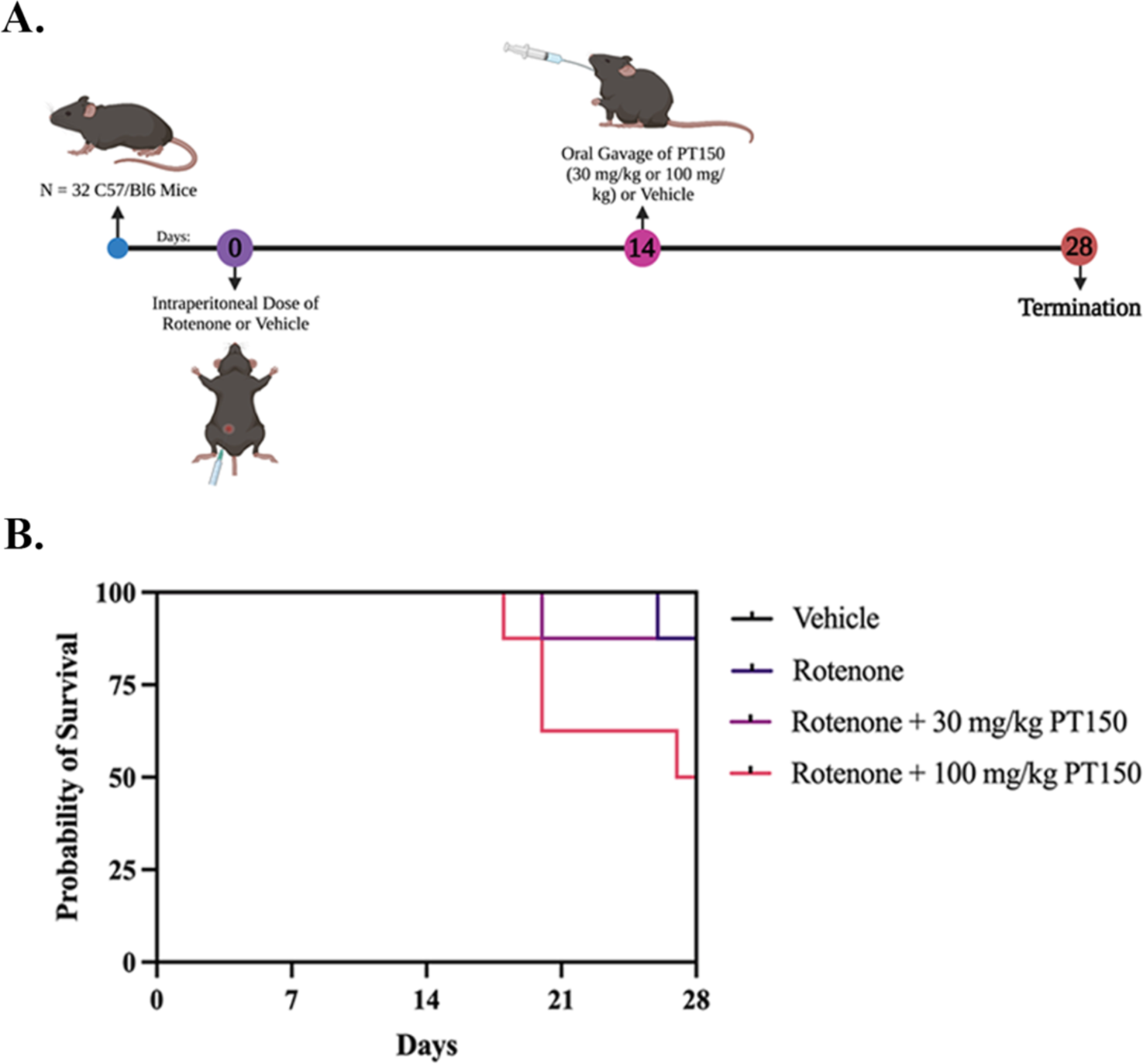
Treatment paradigm and Kaplan-Meier survival curves for each study group. (A) C57BL/6 mice were treated with 2.5 mg/kg/day rotenone for 14 days, immediately followed by treatment with either 30 mg/kg/day or 100 mg/kg/day PT150 by oral gavage for 14 days. Groups: Vehicle control, rotenone, rotenone + 30 mg/kg PT150, rotenone + 100 mg/kg PT150. (B) Survival was reduced in animals treated with 100 mg/kg/day PT150 compared to the other groups. *N* = 8 mice/group.

### PT150 treatment reduces the loss of dopaminergic neurons in the substantia nigra caused by rotenone neurotoxicity

To determine the efficacy of PT150 in treating PD, the extent of neurodegeneration in the SNpc and ST were evaluated. The number of DAn in the SNpc were determined by quantitative stereology of whole-brain tissue sections stained for tyrosine hydroxylase (TH) and neuronal nuclei (NeuN) as previously described [35]. Representative images of DAn in the SNpc of control, rotenone-exposed, and 30 mg/kg PT150-treated animals are shown (**Figure 3A, B, G, H, L**, and **M**). Compared to vehicle controls, animals exposed to rotenone showed a significant decrease, an approximate 50% loss, of TH^+^NeuN^+^ DAn cell bodies in the SNpc by 4 weeks post exposure (**Figure 3F**). Animals exposed to rotenone followed by treatment with 30 mg/kg PT150 had more TH^+^NeuN^+^ cell bodies, which were not statistically different than controls (**Figure 3F**). These data are supported by hematoxylin and eosin staining of brain tissue, which allows for morphological characterization of DAn in the SNpc. In correlation with reduced TH^+^NeuN^+^ cell number, rotenone-exposed animals show cells with pyknotic nuclei (**Figure 3I**, red arrows), whereas PT150-treated animals (**Figure 3N**) appear to have healthy nuclei similar to those observed in the control group (**Figure 3C**, white arrows).

**Figure 3.**
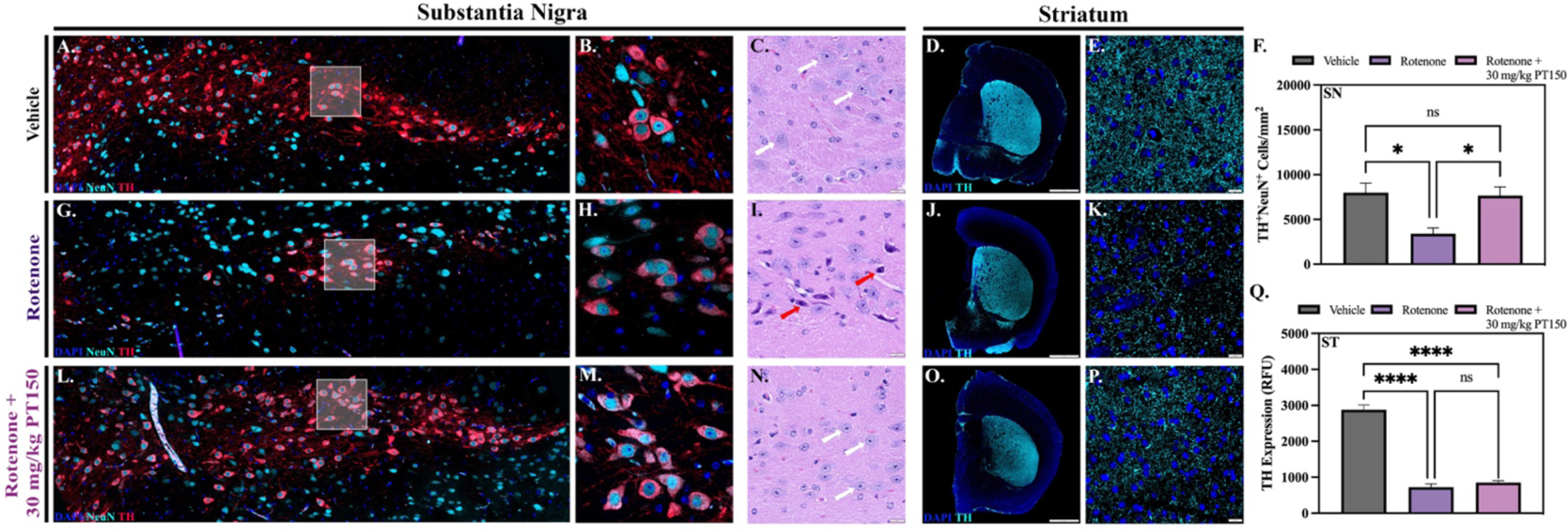
Stereologic and morphologic characterization of dopaminergic neurons in the nigrostriatal system was determined following exposure to rotenone with and without PT150 treatment. Representative images are shown for the substantia nigra (A, B, G, H, L, and M) with immunostaining for dopaminergic neurons (TH, red) and total neurons (NeuN, cyan). The quantity of dopaminergic neuron cell bodies was determined (F). Intensity measurements for TH^+^ projecting fibers in the striatum were performed (Q); representative images of immunostaining for dopaminergic neurons (TH, cyan) are shown (D, E, J, K, O, and P). Parametric one-way ANOVA analysis performed. *N* = 3 – 5/group. \**p* < 0.05; ****p < .0001.

The integrity of DAn axons projecting to the ST was determined by analyzing the fluorescence intensity of TH^+^ immunolabeling in this region. Representative images of control, rotenone-exposed, and 30 mg/kg PT150-treated groups are depicted for the ST (**Figure 3D, E, J, K, O**, and **P**). Striatal DAn terminal integrity was reduced 4 weeks post-exposure to rotenone, as demonstrated by loss of TH^+^ immunostaining intensity compared to vehicle controls (**Figure 3Q**). No protection against loss of dopaminergic terminals was detected in the ST in the PT150 group, as TH^+^ immunostaining in this group was also significantly reduced in the ST compared to controls, and no difference was observed when compared to the rotenone-exposed group (**Figure 3Q**).

### Rotenone-induced microgliosis is reduced in the substantia nigra following PT150 treatment

Microglial reactivity was assessed through quantification of Iba-1^+^ cells in combination with analysis of cellular morphology using immunofluorescence microscopy in the SN and ST of control, rotenone-exposed, and rotenone + 30 mg/kg PT150-treated animals (**Figure 4**). Representative images of Iba-1^+^ immunostaining in the SN (**Figure 4A, F**, and **K**) and ST (**Figure 4B, G**, and **L**) are shown. An increase in the number of Iba-1^+^ cells occurred in the SNpc in response to rotenone exposure, which was significantly reduced by oral treatment with PT150 (**Figure 4C**). Interestingly, no significant difference in the quantity of Iba-1^+^ cells occurred in the SNpr or ST, regardless of experimental group (**Figure 4D** and **E**, respectively).

**Figure 4.**
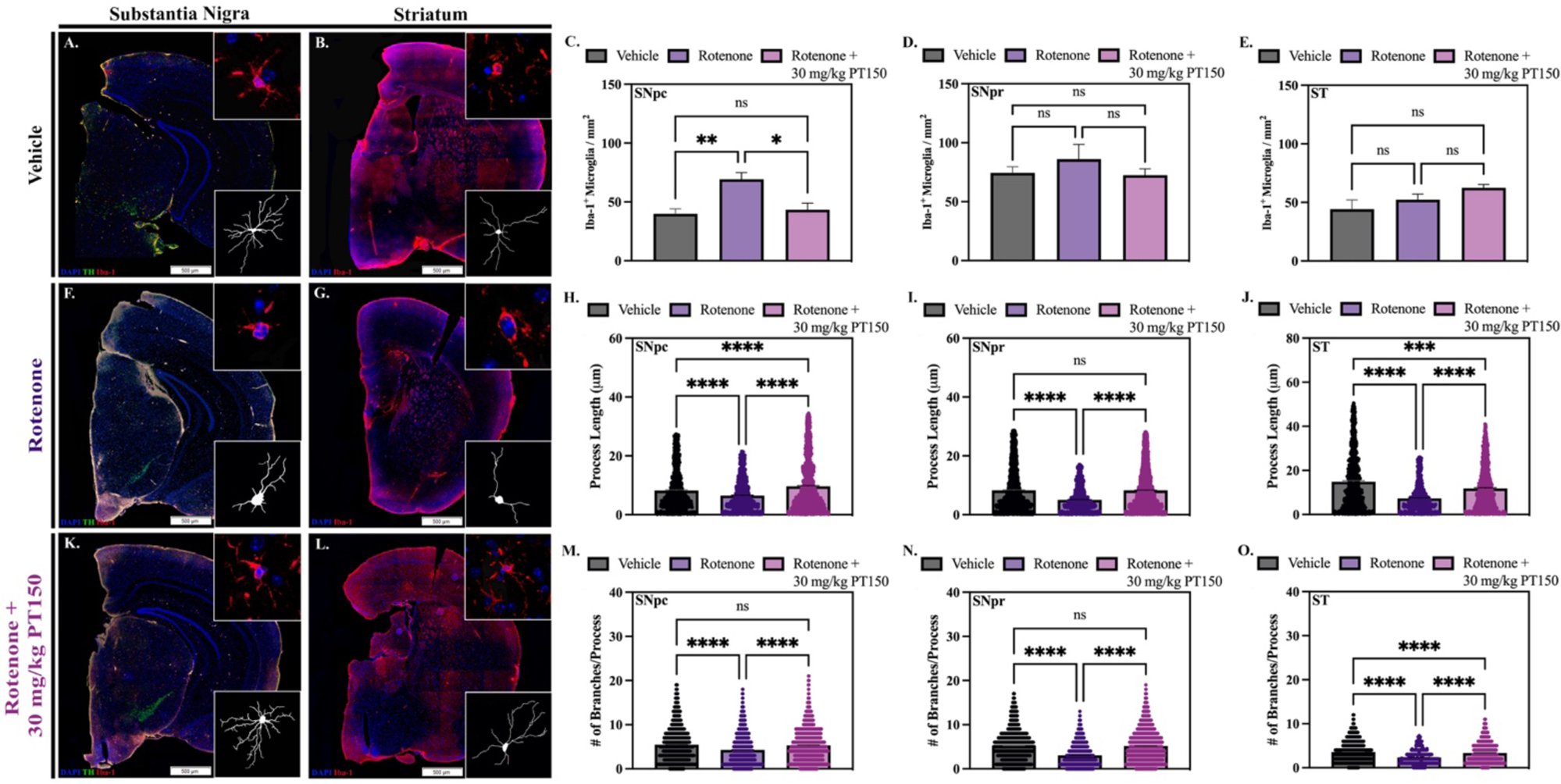
Quantity and morphology of microglia was determined in vehicle, rotenone-exposed, and rotenone + 30 mg/kg PT150-treated animals. Immunofluorescent labeling of microglia (Iba-1, red) was performed; representative montage and high-magnification images are shown for the SN (A, F, and K) and ST (B, G, and L). Analysis of cell quantity was performed using CellSens (C – E) and Imaris software for quantifications of process length (H – J) and the number of process branches (M – O); skeletonized renderings are shown for the SN (A, F, and K) and the ST (B, G, and L). Nonparametric one-way ANOVA with Dunn’s correction analysis performed. *N* = 3 – 5/group. ns = not significant; \**p* < .05; \*\**p* < 0.01; \*\*\**p* < 0.001; \*\*\*\**p* <0 .0001.

Imaris software was used to skeletonize microglial cells in the SN and ST to determine if there were morphological changes between control, rotenone-exposed, and 30 mg/kg PT150-treated groups consistent with varying stages of activation. Skeletonized representations of cells for the SN (**Figure 4A, F**, and **K**) and ST (**Figure 4B, G**, and **L**) are shown for each brain region. Alterations in cellular process length and the number of process branches were quantified as a morphological indication of microglial reactivity. Cellular complexity was reduced in rotenone-exposed animals compared to vehicle controls in the SNpc (**Figure 4H** and **M**), SNpr (**Figure 4I** and **N**), and ST (**Figure 4J** and **O**). Treatment with PT150 significantly increased both length and branching of microglial processes in all three anatomical regions. These data show that treatment with PT150 decreased reactive morphological changes in microglia, but not the overall cell quantity.

### PT150 treatment modulates rotenone-induced astrogliosis in the substantia nigra and the striatum

Reactive astrogliosis was assessed by quantifying the number of astrocytes as well as their secretory and morphometric phenotype. Whole-brain scanning microscopy was used to quantify the number of S100β^+^ cells and GFAP^+^ area, as reactive astrocytes are proliferative and upregulate GFAP. Representative images of control, rotenone-exposed, and rotenone-exposed animals treated with 30 mg/kg PT150 are shown for the SN (**Figure 5A, C**, and **E**) and ST (**Figure 5B, D**, and **F**). Interestingly, no significant difference in the number of S100β^+^ cells is observed following rotenone exposure in any brain region (**Figure 5G, H**, and **I**). A trending decrease in the number of S100β^+^ cells was detected in the SNpc and ST following treatment with PT150, but there were no differences in S100β^+^ cell numbers in the SNpr. Similarly, there was no difference in GFAP^+^ area observed between groups in the SNpc and ST, although PT150 treatment slightly increased GFAP^+^ area compared to controls in the SNpr (**Figure 5J, K**, and **L**).

**Figure 5.**
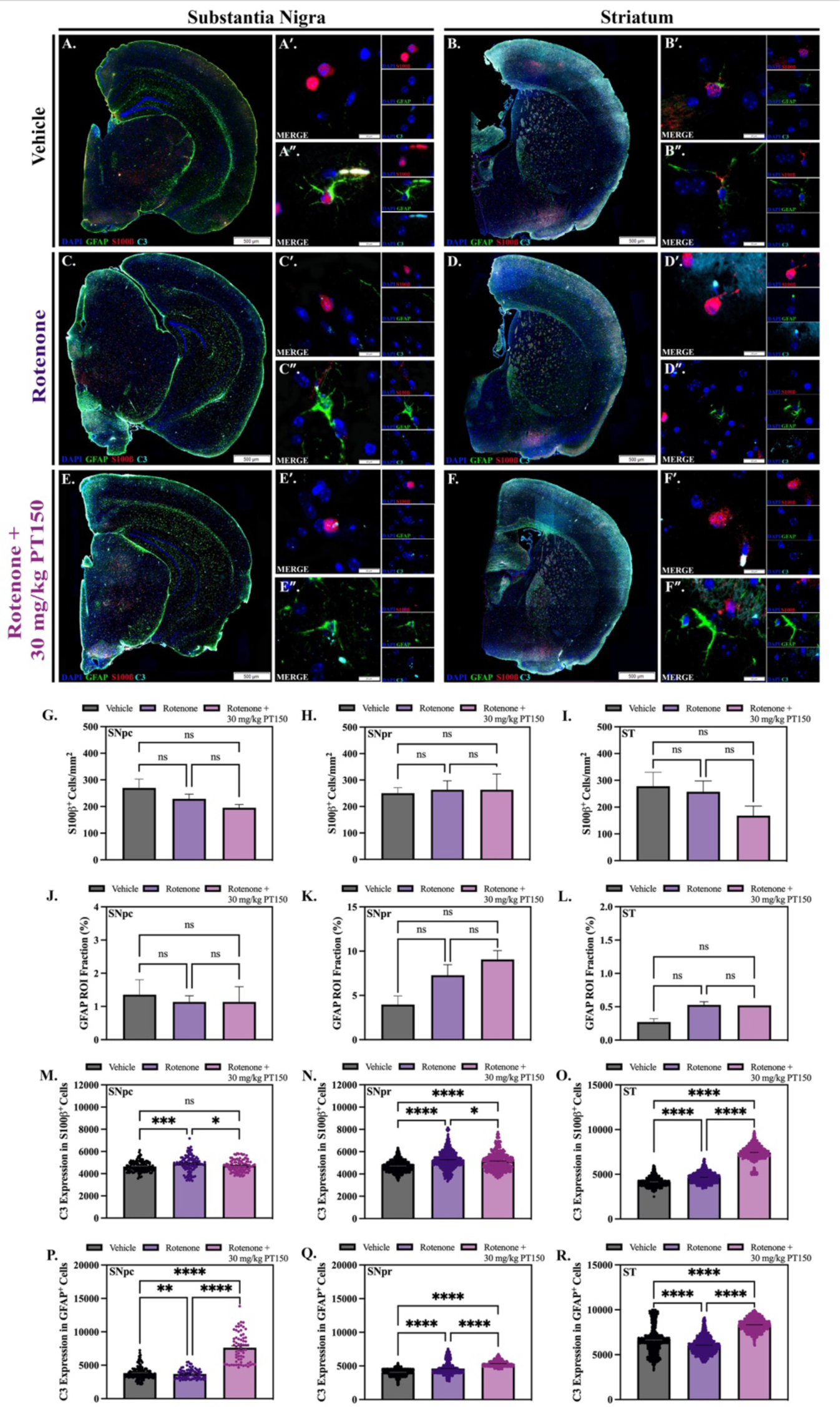
Differential effects of rotenone and PT150 on astrogliosis. Number of astrocytes were quantified in vehicle, rotenone-exposed, and rotenone + 30 mg/kg PT150-treated animals. Immunofluorescent labeling of astrocytes (GFAP, green; S100β, red; C3, cyan) was performed; representative montage and high-magnification images are shown for the SN (A, C, and E) and ST (B, D, and F). The quantity of S100β^+^ cells is shown for the SNpc (G), SNpr (H), and ST (I). Immunostaining was quantified for the maximum expression of GFAP (% per regional area) in the SNpc (J), SNpr (K), and ST (L). Maximal C3 expression in S100β^+^ soma and GFAP^+^ processes were determined within the SNpc (M, P), SNpr (N, Q), and ST (O, R), respectively. Nonparametric one-way ANOVA with Dunn’s correction analysis performed. *N* = 3 – 5/group. ns = not significant; \**p* < 0.05; \*\**p* <0 .01; \*\*\**p* < 0.001; \*\*\*\**p* <0 .0001.

High-resolution fluorescence microscopy was also used to evaluate the phenotype of astrocytes by examining the cellular distribution of C3 in S100β^+^ cell soma and GFAP^+^ processes as an indication of inflammatory activation. The co-expression of C3 and either GFAP or S100β was quantitively analyzed in the SNpc, SNpr, and ST (**Figure 5**). Representative images of control, rotenone-exposed, and rotenone-exposed animals treated with 30 mg/kg PT150 are shown for the SN (**Figure 5A, C**, and **E**) and ST (**Figure 5B, D**, and **F**). Expression of C3 within astrocyte soma in the SNpc was significantly increased following rotenone exposure, but decreased with PT150 treatment, where no significant difference between control and PT150-treated animals was observed (**Figure 5M**). C3 in astrocyte processes in PT150-treated animals was significantly increased compared to both control and rotenone-exposed animals in this same brain region (**Figure 5P**). In the SNpr, somal expression of C3 was significantly increased in PT150-treated animals compared to controls but was not as high as the rotenone-exposed group (**Figure 5N**). Expression of C3 in processes was upregulated following both rotenone exposure and PT150 treatment in this anatomical region (**Figure 5Q**). Interestingly, C3 expression in both the soma and processes was significantly higher in PT150-treated animals compared to the other groups in the ST (**Figure 5O** and **R**).Morphological changes of the astrocytes, including process length, area, and branching, were analyzed using Imaris software; representative images of control (**Figure 6A** and **B**), rotenone-exposed (**Figure 6C** and **D**), and rotenone-exposed animals treated with 30 mg/kg PT150 (**Figure 6E** and **F**) are shown for the SN and ST, respectively. In the SNpc, there was no significant difference in cell body size between groups, as determined by S100β^+^ cell area (**Figure 6G**). Exposure to rotenone resulted in astrocytes with decreased process branching (**Figure 6P**) and length (**Figure 6M**), but no change in process area (**Figure 6J**), compared to controls in this same brain region. PT150 treatment further decreased process branching (**Figure 6P**) and length (**Figure 6M**), but increased process area (**Figure 6J**) compared to vehicle controls in the SNpc. In the SNpr, significant cell body hypertrophy is demonstrated in animals exposed to rotenone. This morphologic response was increased in animals treated with PT150 compared to controls but was not different from rotenone-treated animals (**Figure 6H**). Rotenone-exposed animals contain astrocytes with morphology similar to controls, with no significant difference in process area (**Figure 6K**) or length (**Figure 6N**), but an increase in branching (**Figure 6Q**). PT150 treatment significantly increased process branching (**Figure 6Q**) and area (**Figure 6K**), but not length (**Figure 6N**). In the ST, no evidence of hypertrophic cell body changes were observed (**Figure 6I**), although both rotenone exposure and PT150 treatment resulted in process elongation (**Figure 6O**) and increased branching (**Figure 6R**) compared to controls. Altogether, these data establish that PT150 treatment modulated astrocyte reactivity towards a more activated morphological phenotype in a brain region-dependent manner.

**Figure 6.**
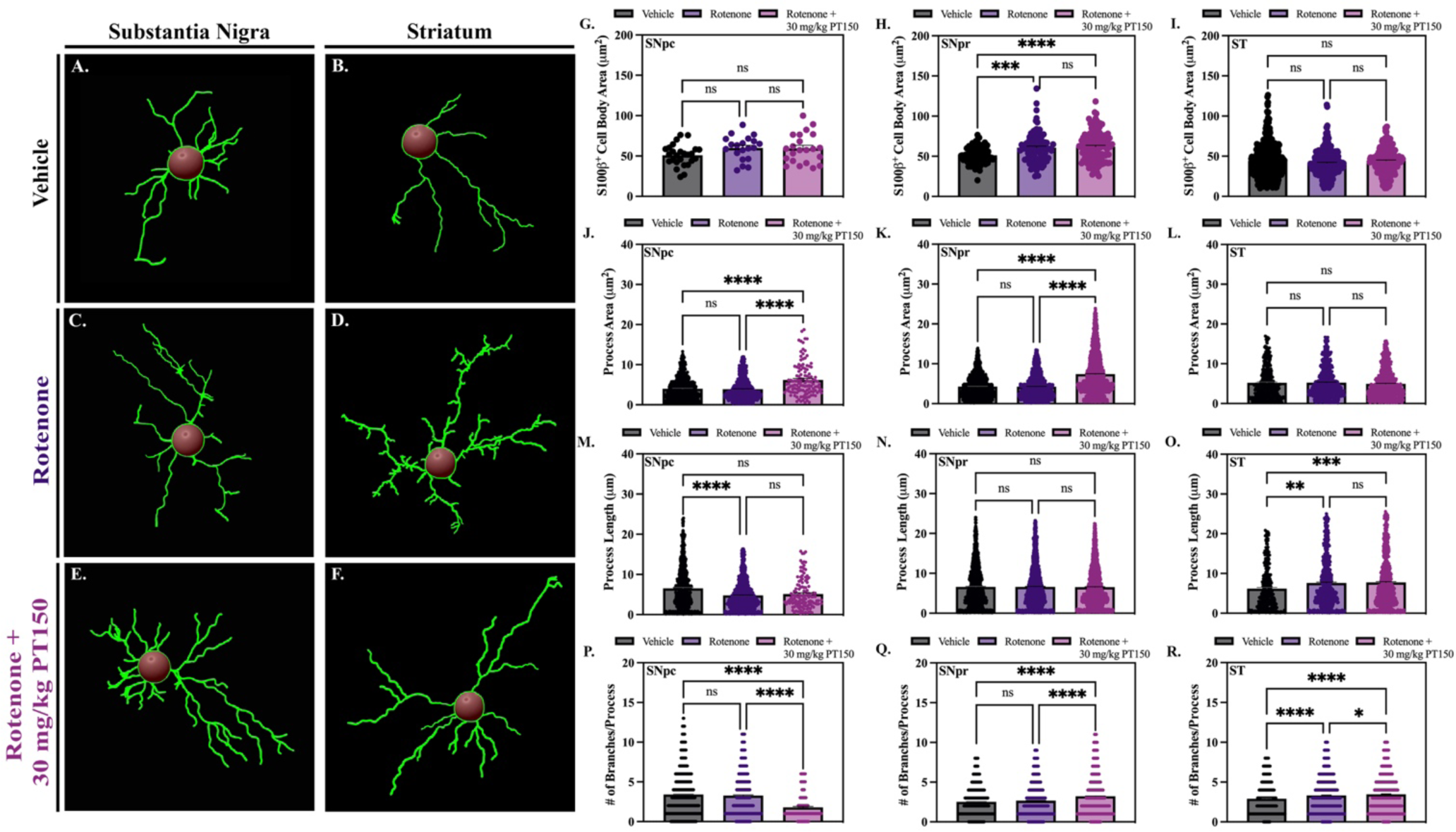
Astrocyte morphology was evaluated in vehicle, rotenone-exposed, and rotenone + 30 mg/kg PT150-treated animals. Morphological analysis was performed using CellSens for quantification of cell body area (G – I) and Imaris software for quantifications of process area (J – L), process length (M – O) and the number of process branches (P – R); skeletonized renderings are shown for the SN (A, C, and E) and the ST (B, D, and F). Nonparametric one-way ANOVA with Dunn’s correction analysis performed. *N* = 3 – 5/group. ns = not significant; \*\**p* < 0.05; \*\**p* <0 .01; \*\*\**p* < 0.001; \*\*\*\**p* <0 .0001.

### PT150 treatment reduces accumulation of α-synuclein in neurons and alters glial trafficking of phosphorylated α-synuclein

Accumulation of α-synuclein was determined by immunofluorescence staining for the aggregation-prone phospho-serine 129 form of the protein (p129 α-syn) in the SN (**Figure 7**). The extent of accumulation of misfolded protein in neurons was determined by analyzing the intensity of p129 α-syn in TH^+^ neurons in the SNpc. Representative images of control, rotenone-exposed, and rotenone + 30 mg/kg PT150-treated animals are shown (**Figure 7, D**, and **G**). Rotenone-exposed animals had an increase in intra-neuronal p129 α-syn compared to control animals (**Figure 7J**). Notably, p129 α-syn expression was reduced in DAn in the PT150-treated animals (**Figure 7J**).

**Figure 7.**
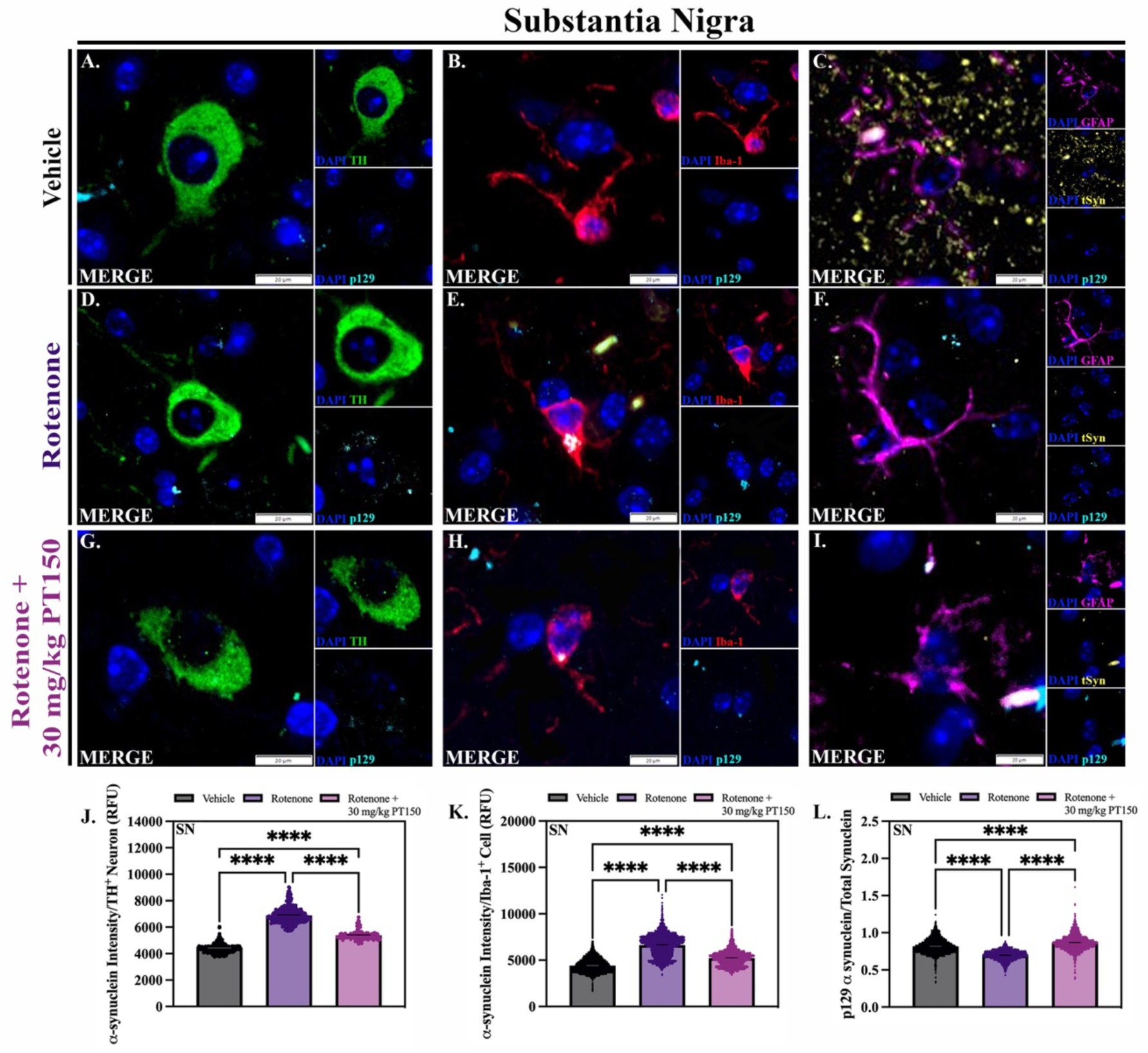
Accumulation and trafficking of α-synuclein. Levels of α-synuclein were evaluated in dopaminergic neurons, microglia, and astrocytes in vehicle, rotenone-exposed, and rotenone + 30 mg/kg PT150-treated animals. Immunofluorescent labeling of α-synuclein phosphorylated at serine 129 (p129, cyan), dopaminergic neurons (TH, green), microglia (Iba-1, red), and astrocytes (GFAP, purple) was performed; representative high-magnification images are shown for the SN. Maximal p129 expression within dopaminergic neurons (J), microglia (K), and astrocytes (L) was determined. Nonparametric one-way ANOVA with Dunn’s correction analysis performed. *N* = 3 – 5/group. \*\*\*\**p* < 0.0001.

To determine the role glia may play in removing or trafficking misfolded α-synuclein in the SN, p129 α-syn accumulation was also analyzed within GFAP^+^ astrocytes and Iba-1^+^ microglia. There is a significant increase in p129 α-syn within Iba-1 cells following rotenone exposure (**Figure 7K**). Animals treated with PT150 show more p129 α-syn within microglia compared to controls, but significantly less than the untreated, rotenone-exposed group (**Figure 7K**). Although less p129 α-syn was detected in GFAP^+^ cells in both the rotenone-exposed and PT150-treated groups (**Supplemental Figure 1**), ratiometric analysis of p129 α-syn expression to total synuclein shows that PT150 treatment drives removal of p129 α-syn in the SN compared to astrocytes in the rotenone-exposed group (**Figure 7L, Supplemental Figure 1**).

## Discussion

With cases of PD projected to double over the next twenty years, it is imperative that therapeutic options be developed to improve the quality of life for those who are affected [38]. Current dopamine mimetics treat the motor symptoms of PD but are ineffective at slowing the progression of disease. Here, we report that post-lesioning treatment with the GR modulator, PT150, reduced loss of DAn in the SNpc of C57Bl/6 mice in the rotenone model of PD. Compared with other models of PD, the rotenone model accurately recapitulates many of the key pathological hallmarks of the disease, including inhibition of mitochondrial complex I, degeneration of DAn, and the formation of intraneuronal Lewy bodies [6]. In parallel to the neuropathological progression of clinical disease, rotenone neurotoxicity in murine models results in glial reactivity peaking at 2 weeks post-exposure, followed by synucleinopathy and DAn loss in the nigrostriatal pathway by 4 weeks post-exposure [7]. Computational-based molecular docking studies indicate that PT150 preferentially binds to and interacts with the co-activator site of the ligand binding domain of GR rather than the steroid binding site, due to a lower steric and conformationally restricted penalty for binding (**Figure 1**). Molecular docking studies demonstrated that PT150 has the potential to bind the co-activator peptide site, and it is logical to assume that this site would be more accessible compared with the steroid binding site based on restricted access to the steroid binding pocket in the intact protein structure. PT150 interacts favorably with the co-activator site, which would induce a conformational change in the receptor upon binding the co-activator domain, suggesting this as a site of allosteric modulation that could inhibit transcriptional activity without directly competing for binding with endogenous corticosteroid ligands. Due to allosteric modulation of GR by PT150, we postulated that pharmacologic modulation of this receptor would be neuroprotective by reducing pro-inflammatory activity amongst microglia and astrocytes, as well as preventing the propagation of misfolded α-syn in neurons. In the present study, mice were exposed to daily intraperitoneal administration of rotenone or vehicle for 14 days, followed by treatment with vehicle, 30 mg/kg, or 100 mg/kg PT150 by oral gavage for 14 days (**Figure 2**). Mortality occurred in approximately 50% of the animals exposed to 100 mg/kg PT150, whereas 30 mg/kg was well-tolerated, as demonstrated by a mortality rate similar to the rotenone-exposed group (**Figure 2**).

Oral treatment with PT150 immediately following daily rotenone exposure preserved the number of TH^+^ neuronal cell bodies in the SN, although no protection of striatal axons was seen (**Figure 3**). Although incompletely understood, it is thought that axonal degeneration occurs prior to complete loss of DAn cell bodies and is the primary determinant of clinical disease [39, 40]. Therefore, because PT150 treatments began after exposure to rotenone, it is likely that the majority of degeneration of DAn axons had occurred prior to therapeutic intervention. Although decreased expression of TH is often correlated to neurodegeneration, altered expression of intracellular enzymes may be independent of neuronal survival following stress or neurotoxic injury [41]. Acute damage mediated by rotenone may result in fluctuations in enzymatic expression in terminals, and subsequently synaptic dysfunction, but TH immunostaining does not definitively indicate complete neuronal degeneration in the ST [42]. Additionally, though loss of cell bodies is irreversible, neurons within the CNS have, to a certain extent, the capacity to induce regrowth of axons and rescue function following injury. This suggests that the disruptions observed in the ST following PT150 treatment may not have permanent clinical implications. Ultimately, PT150 treatment was neuroprotective, as it was effective at preventing the irreversible loss of DAn cell bodies in the SNpc following rotenone neurotoxicity.

Neuroinflammation is an early critical feature of PD, and DAn are especially vulnerable to the toxic effects of pro-inflammatory mediators and subsequent oxidative stress [43]. Therefore, inhibiting neuroinflammatory signaling has therapeutic potential [44, 45]. The ability of the GR to modulate key inflammatory pathways identified in PD, as well as the high quantities of the receptor identified on both DAn and glia in rodent models, suggests that targeting the GR is a promising avenue for treating PD. Although largely anti-inflammatory effects of GR modulation have been described, it must be noted that GCs do not have solely immunosuppressive actions. GR agonism can sometimes increase pro-inflammatory responses in the brain, though these inflammatory actions have, thus far, been described primarily in the hippocampus [46]. The ubiquitous expression of GRs within parenchymal cells of the CNS also means there is the potential for undesired effects in some cell populations, such that therapeutics that solely act as GR agonists or antagonists may prove ineffective or problematic. This is demonstrated by other models of PD, such as 1-methyl-4-phenyl-1,2,3,6-tetrahydropyridine (MPTP) neurotoxicity, where inactivating astrocytic GRs exacerbated DAn loss [21, 46]. Because PT150 is thought to allosterically modulate GR by binding to the co-activator domain of the receptor and thereby inhibit pro-inflammatory transcription, it may function as a better inhibitor of neuroinflammation [25]. In support of this mechanism, we observed decreased quantities of reactive microglia in the SNpc in PT150-treated animals (**Figure 4**). A reduction in microgliosis was not observed in the ST of animals treated with PT150, as no change in cell quantity was observed, although the cells in this region had an anti-inflammatory or inactivated morphology (**Figure 4**). Reactive, or disease-associated, microglia (DAM’s) are described as having an ameboid-like morphology, with decreased branch length and volume, as well as increased cell body size [47–49]. They readily produce reactive species and pro-inflammatory mediators, which, when prolonged, contributes to the progression of neuronal injury and the misfolding of native proteins into neurotoxic forms [50, 51]. Dystrophic microglia have been implicated in both clinical and laboratory models of disease, and sustained microgliosis has been identified in human PD patients [7, 52]. In the present study, the observed reduction in both the number and reactivity of microglia in the SNpc indicates that PT150 treatment reduces pro-inflammatory microglial responses, but this occurs in a brain region-dependent manner. The suppressed microglial response following PT150 treatment in the SN may be beneficial, as experimental evidence from others demonstrates that reducing microgliosis can protect DAn [22].

Although increased numbers of reactive glial cells have been described in numerous neurodegenerative diseases, the function of activated astrocytes cannot be grouped simply as neurotoxic or neuroprotective, but instead resides within a spectrum of molecular and cellular changes that dictate their function [53, 54]. Chronic or severe neuroinflammatory activation of astrocytes contributes to neuronal injury, but complete inhibition of glial activation can lead to pathogenic protein accumulation and the loss of critical homeostatic functions, like neurotransmitter regulation and cellular repair mechanisms [55]. The extent of astrogliosis was evaluated through analysis of the quantity, morphology, and secretory phenotype of the astrocytes present in the SN and ST, including C3^+^ immunolabeling in the cell body (S100β) and processes (GFAP) to identify reactive astrocytes. Morphological measures included cell body area, the number of branches per process, process length, and process area. Upon activation, astrocytes increase process complexity and volume, as well as elongate them, which allows these cells to facilitate cellular communication [56]. Activated cells upregulate production of complement in an effort to opsonize pathogens and recruit immune cells, but overactivation of the complement pathway can contribute to neuronal damage and overpruning of synapses [57]. In the present study, rotenone exposure induced activated morphology and robust production of C3 by astrocytes in the SN and ST (**Figure 5** and **6**). Unexpectedly, these cellular responses were augmented following PT150 treatment. An increase in C3 is identified in the cell bodies and processes of astrocytes in both the SN and ST following PT150 exposure (**Figure 5**). The astrocytes in these regions also increase process length and branching and demonstrate cell body hypertrophy in a brain-region dependent manner (**Figure 6**).

This apparent increase in glial reactivity may have neuroprotective functions. More complex and longer processes on activated astrocytes allow for increased contact with nearby cells and vasculature, which permits them to stimulate cell-to-cell communication and blood-brain barrier tightness [56]. During disease states, activated astrocytes secrete pro-inflammatory cytokines and other factors, such as glial derived neurotrophic factor (GDNF) and Gata3, that stimulate neurogenesis [55, 58–62]. Some evidence suggests that astrocytes can also facilitate neuronal regeneration through their collagen-rich processes, which form scaffolds for the formation of neurites [63]. Furthermore, while neurotoxic effects of C3 have been described, it’s role in disease is context-dependent, and therefore may be neuroprotective in some situations [64]. Complement is involved in cellular activation and recruitment, thereby increasing microglial responses like phagocytosis, which removes debris and prohibits the formation of misfolded protein aggregates [65]. Activated astrocytes have also been implicated in the removal of cellular debris in the ST, which can decrease microglia-mediated degeneration of DAn in the SN [66]. Models of Alzheimer’s Disease show that C3 deficiency can increase amyloid burden and decrease neuronal viability, highlighting the neuroprotective function of this protein in the brain [67, 68]. It is possible that the morphological and secretory indications of astroglial reactivity documented in response to PT150 treatment stimulates neuroprotective mechanisms, enhancing the capacity of these cells to mitigate damage caused by microglial populations and produce factors that stimulate neuronal regeneration.

Another key pathological feature of PD is the aggregation of hyperphosphorylated α-syn in DAn cell bodies of the SNpc. This neurotoxic form of synuclein mediates synaptic disturbances, mitochondrial and lysosomal dysfunction, and persistent neuroinflammatory signaling, ultimately contributing to neuronal death [69]. Neurons secrete soluble forms of α-syn in an effort to reduce toxicity, that is then endocytosed by nearby cells. This process allows for cell-to-cell spread of the protein in a prion-like manner that results in insoluble aggregates throughout the brain [70–72]. Glia surrounding DAn phagocytose and degrade secreted α-syn, thereby clearing neurotoxic proteins from the brain. Following PT150 treatment, a reduction in the accumulation of phosphorylated α-syn was observed in DAn in the SN compared to rotenone-exposed, untreated animals (**Figure 7**). A decrease in α-syn within Iba-1^+^ cells also occurred following treatment, suggesting that the microglia in the SN, which have an anti-inflammatory morphology, may reduce phagocytosis of α-syn (**Figure 7**). Interestingly, when evaluating the ratio of phosphorylated α-syn to total synuclein within astrocytes, an increase in observed (**Figure 7**, **Supplemental Figure 1**). This suggests that the astrocytes in the SN, which show markers of having an activated phenotype, may be upregulating their capacity to phagocytose α-syn. Although the state of activation in which astrocytes enhance their phagocytic capacity is sometimes contradictory, such a mechanism has been established in reactive astrocytes identified in other laboratory models [73, 74]. This function allows astrocytes to efficiently remove fibrillar α-syn through lysosomal degradation, decreasing the pathogenic accumulation of α-syn in neurons [75]. Although reducing glial responses may be neuroprotective, it can also perpetuate synucleinopathy. It is possible that through modulation of glia by PT150, astrocytes in the SN increase their capacity for removal of neurotoxic α-syn, thereby promoting survival of DAn in the SN.

Collectively, this study highlights the potential role of GR modulation as a method of interfering with the progression of PD neuropathology. We demonstrate that treatment with 30 mg/kg of the novel therapeutic PT150 following exposure to rotenone reduced microglial reactivity, modulated the astrocytic response, and interfered with the accumulation of hyperphosphorylated α-syn in DAn. Overall, PT150 reduced the loss of DAn cell bodies in the SN, although was not completely protective, as demonstrated by evidence of axonal dysfunction in the ST. Altogether, these findings effectively illuminate neuroprotective mechanisms of targeting the GR to impede PD neuropathology.

## Acknowledgements

The authors thank Dr. Neil Theise (Chief Scientific Officer, Pop Test Therapeutics, LLC) for providing PT150. This work was supported by NIH-ES035043 and DOD-W81XWH-22-1-0663 (to RBT).

---

**Supplemental Figure 1.**
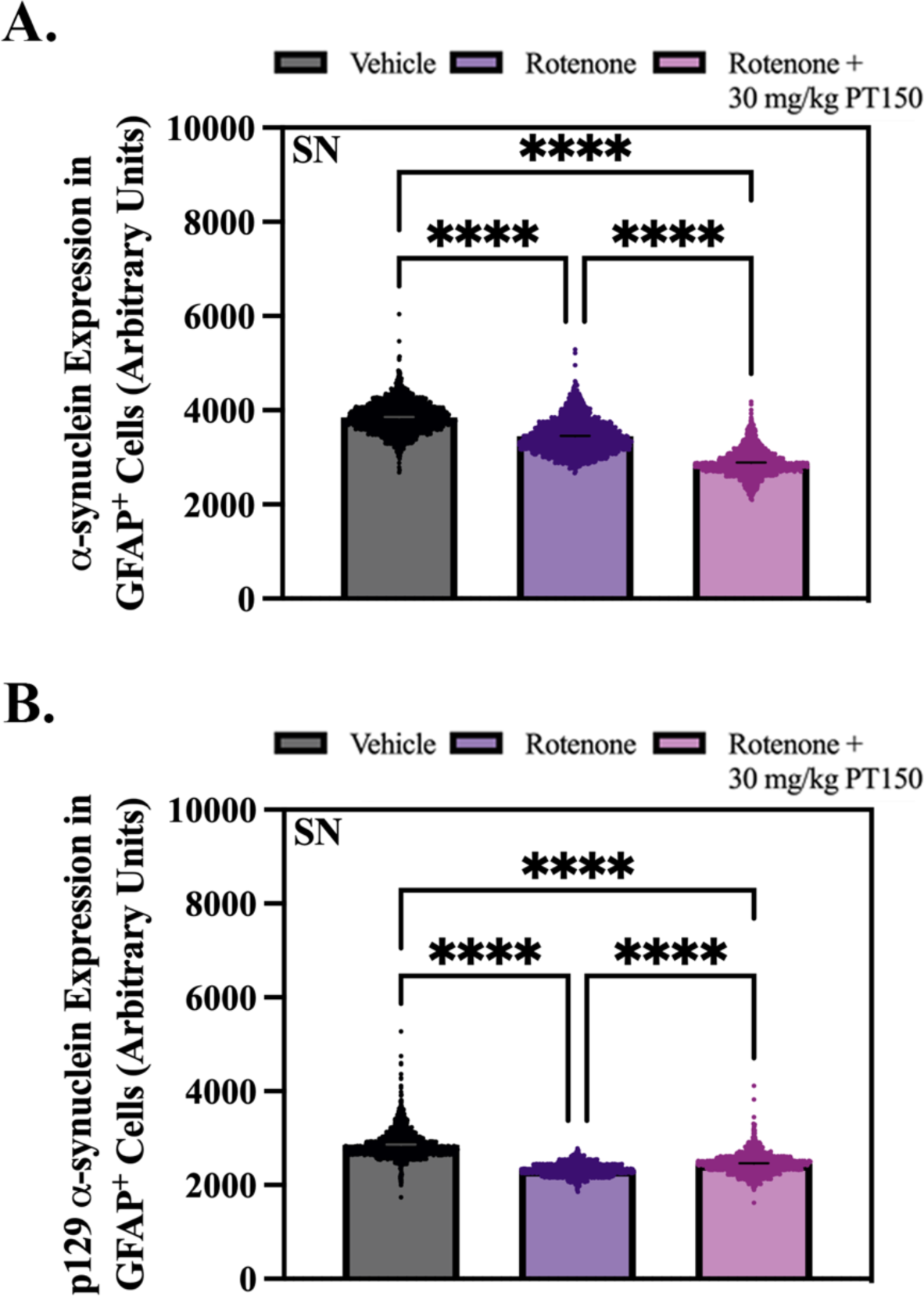
Analysis of phosphorylated and total α-synuclein within GFAP^+^ astrocytes was performed using immunofluorescence microscopy. Intracellular expression of α-synuclein (A) and α-synuclein phosphorylated at serine 129 (B) is shown. Nonparametric one-way ANOVA with Dunn’s correction analysis performed. *N* = 3 – 5/group. \*\*\*\**p* < 0.0001.

## References

1. Global, regional, and national burden of neurological disorders, 1990-2016: a systematic analysis for the Global Burden of Disease Study 2016. Lancet Neurol, 2019. 18(5): p. 459–480.

2. Poewe, W., et al., Parkinson disease. Nature Reviews Disease Primers, 2017. 3(1): p. 17013.

3. Willis, A.W., et al., Incidence of Parkinson disease in North America. npj Parkinson’s Disease, 2022. 8(1): p. 170.

4. Sveinbjornsdottir, S., The clinical symptoms of Parkinson’s disease. J Neurochem, 2016. 139 Suppl 1: p. 318–324.

5. Tanner, C.M., et al., Rotenone, paraquat, and Parkinson’s disease. Environ Health Perspect, 2011. 119(6): p. 866–72.

6. Li, N., et al., Mitochondrial complex I inhibitor rotenone induces apoptosis through enhancing mitochondrial reactive oxygen species production. J Biol Chem, 2003. 278(10): p. 8516–25.

7. Rocha, S.M., et al., Rotenone induces regionally distinct α-synuclein protein aggregation and activation of glia prior to loss of dopaminergic neurons in C57Bl/6 mice. Neurobiol Dis, 2022. 167: p. 105685.

8. Sheeler, C., et al., Glia in Neurodegeneration: The Housekeeper, the Defender and the Perpetrator. Int J Mol Sci, 2020. 21(23).

9. Kwon, H.S. and S.-H. Koh, Neuroinflammation in neurodegenerative disorders: the roles of microglia and astrocytes. Translational Neurodegeneration, 2020. 9(1): p. 42.

10. Lema Tomé, C.M., et al., Inflammation and α-synuclein’s prion-like behavior in Parkinson’s disease--is there a link? Mol Neurobiol, 2013. 47(2): p. 561–74.

11. Tansey, M.G., M.K. McCoy, and T.C. Frank-Cannon, Neuroinflammatory mechanisms in Parkinson’s disease: Potential environmental triggers, pathways, and targets for early therapeutic intervention. Experimental Neurology, 2007. 208(1): p. 1–25.

12. Hensleigh, E. and L.M. Pritchard, Glucocorticoid receptor expression and sub-cellular localization in dopamine neurons of the rat midbrain. Neurosci Lett, 2013. 556: p. 191–5.

13. Barnes, P.J., Anti-inflammatory actions of glucocorticoids: molecular mechanisms. Clin Sci (Lond), 1998. 94(6): p. 557–72.

14. Landfield, P.W., et al., A new glucocorticoid hypothesis of brain aging: implications for Alzheimer’s disease. Curr Alzheimer Res, 2007. 4(2): p. 205–12.

15. Ros-Bernal, F., et al., Microglial glucocorticoid receptors play a pivotal role in regulating dopaminergic neurodegeneration in parkinsonism. Proceedings of the National Academy of Sciences, 2011. 108(16): p. 6632–6637.

16. Perlman, W.R., et al., Age-related differences in glucocorticoid receptor mRNA levels in the human brain. Neurobiology of Aging, 2007. 28(3): p. 447–458.

17. Caudal, D., et al., Depressive-like phenotype induced by AAV-mediated overexpression of human α-synuclein in midbrain dopaminergic neurons. Experimental Neurology, 2015. 273: p. 243–252.

18. Herrero, M.-T., et al., Inflammation in Parkinson’s disease: role of glucocorticoids. Frontiers in Neuroanatomy, 2015. 9.

19. Atbin, D., et al., Salivary cortisol levels in Parkinson&#039;s disease and its correlation to risk behaviour. Journal of Neurology, Neurosurgery &amp;amp; Psychiatry, 2011. 82(10): p. 1107.

20. Ouanes, S. and J. Popp, High Cortisol and the Risk of Dementia and Alzheimer’s Disease: A Review of the Literature. Frontiers in Aging Neuroscience, 2019. 11.

21. Maatouk, L., et al., Glucocorticoid receptor in astrocytes regulates midbrain dopamine neurodegeneration through connexin hemichannel activity. Cell Death & Differentiation, 2019. 26(3): p. 580–596.

22. Rocha, S.M., et al., Microglia-specific knock-out of NF-κB/IKK2 increases the accumulation of misfolded α-synuclein through the inhibition of p62/sequestosome-1-dependent autophagy in the rotenone model of Parkinson’s disease. Glia, 2023. 71(9): p. 2154–2179.

23. Le, W., J. Wu, and Y. Tang, Protective Microglia and Their Regulation in Parkinson’s Disease. Front Mol Neurosci, 2016. 9: p. 89.

24. Morice, C., et al., A randomized trial of safety and pharmacodynamic interactions between a selective glucocorticoid receptor antagonist, PT150, and ethanol in healthy volunteers. Scientific Reports, 2021. 11(1): p. 9876.

25. Rocha, S.M., et al., A Novel Glucocorticoid and Androgen Receptor Modulator Reduces Viral Entry and Innate Immune Inflammatory Responses in the Syrian Hamster Model of SARS-CoV-2 Infection. Frontiers in Immunology, 2022. 13.

26. Biggadike, K., et al., X-ray crystal structure of the novel enhanced-affinity glucocorticoid agonist fluticasone furoate in the glucocorticoid receptor-ligand binding domain. J Med Chem, 2008. 51(12): p. 3349–52.

27. Kauppi, B., et al., The three-dimensional structures of antagonistic and agonistic forms of the glucocorticoid receptor ligand-binding domain: RU-486 induces a transconformation that leads to active antagonism. J Biol Chem, 2003. 278(25): p. 22748–54.

28. Friesner, R.A., et al., Glide: a new approach for rapid, accurate docking and scoring. 1. Method and assessment of docking accuracy. J Med Chem, 2004. 47(7): p. 1739–49.

29. Friesner, R.A., et al., Extra precision glide: docking and scoring incorporating a model of hydrophobic enclosure for protein-ligand complexes. J Med Chem, 2006. 49(21): p. 6177–96.

30. Halgren, T.A., Identifying and characterizing binding sites and assessing druggability. J Chem Inf Model, 2009. 49(2): p. 377–89.

31. Beckstein, O., A. Fourrier, and B.I. Iorga, Prediction of hydration free energies for the SAMPL4 diverse set of compounds using molecular dynamics simulations with the OPLS-AA force field. J Comput Aided Mol Des, 2014. 28(3): p. 265–76.

32. Cannon, J.R., et al., A highly reproducible rotenone model of Parkinson’s disease. Neurobiol Dis, 2009. 34(2): p. 279–90.

33. Reagan-Shaw, S., M. Nihal, and N. Ahmad, Dose translation from animal to human studies revisited. The FASEB Journal, 2008. 22(3): p. 659–661.

34. Hippius, H. and G. Neundörfer, The discovery of Alzheimer’s disease. Dialogues in Clinical Neuroscience, 2003. 5(1): p. 101–108.

35. Tapias, V. and J.T. Greenamyre, A Rapid and Sensitive Automated Image-Based Approach for In Vitro and In Vivo Characterization of Cell Morphology and Quantification of Cell Number and Neurite Architecture. Current Protocols in Cytometry, 2014. 68(1): p. 12.33.1–12.33.22.

36. Bantle, C.M., et al., Infection with mosquito-borne alphavirus induces selective loss of dopaminergic neurons, neuroinflammation and widespread protein aggregation. npj Parkinson’s Disease, 2019. 5(1): p. 20.

37. Vardjan, N., et al., IFN-γ-induced increase in the mobility of MHC class II compartments in astrocytes depends on intermediate filaments. Journal of Neuroinflammation, 2012. 9(1): p. 144.

38. Dorsey, E.R., et al., The Emerging Evidence of the Parkinson Pandemic. J Parkinsons Dis, 2018. 8(s1): p. S3–s8.

39. Salvadores, N., et al., Axonal Degeneration during Aging and Its Functional Role in Neurodegenerative Disorders. Front Neurosci, 2017. 11: p. 451.

40. Tagliaferro, P. and R.E. Burke, Retrograde Axonal Degeneration in Parkinson Disease. Journal of Parkinson’s Disease, 2016. 6: p. 1–15.

41. Lams, B.E., O. Isacson, and M.V. Sofroniew, Loss of transmitter-associated enzyme staining following axotomy does not indicate death of brainstem cholinergic neurons. Brain Research, 1988. 475(2): p. 401–406.

42. Salvatore, M.F., et al., Tyrosine Hydroxylase Inhibition in Substantia Nigra Decreases Movement Frequency. Mol Neurobiol, 2019. 56(4): p. 2728–2740.

43. Ni, A. and C. Ernst, Evidence That Substantia Nigra Pars Compacta Dopaminergic Neurons Are Selectively Vulnerable to Oxidative Stress Because They Are Highly Metabolically Active. Front Cell Neurosci, 2022. 16: p. 826193.

44. Liu, M. and G. Bing, Lipopolysaccharide Animal Models for Parkinson’s Disease. Parkinson’s Disease, 2011. 2011: p. 327089.

45. Surmeier, D.J., Determinants of dopaminergic neuron loss in Parkinson’s disease. Febs j, 2018. 285(19): p. 3657–3668.

46. De Nicola, A.F., et al., Insights into the Therapeutic Potential of Glucocorticoid Receptor Modulators for Neurodegenerative Diseases. Int J Mol Sci, 2020. 21(6).

47. Heindl, S., et al., Automated Morphological Analysis of Microglia After Stroke. Frontiers in Cellular Neuroscience, 2018. 12.

48. Fernández-Arjona, M.d.M., et al., Microglial Morphometric Parameters Correlate With the Expression Level of IL-1β, and Allow Identifying Different Activated Morphotypes. Frontiers in Cellular Neuroscience, 2019. 13.

49. Morrison, H., et al., Quantitative microglia analyses reveal diverse morphologic responses in the rat cortex after diffuse brain injury. Scientific Reports, 2017. 7(1): p. 13211.

50. Simpson, D.S.A. and P.L. Oliver, ROS Generation in Microglia: Understanding Oxidative Stress and Inflammation in Neurodegenerative Disease. Antioxidants (Basel), 2020. 9(8).

51. Smith, J.A., et al., Role of pro-inflammatory cytokines released from microglia in neurodegenerative diseases. Brain Res Bull, 2012. 87(1): p. 10–20.

52. Gerhard, A., et al., In vivo imaging of microglial activation with [11C](R)-PK11195 PET in idiopathic Parkinson’s disease. Neurobiology of Disease, 2006. 21(2): p. 404–412.

53. Liddelow, S.A. and B.A. Barres, Reactive Astrocytes: Production, Function, and Therapeutic Potential. Immunity, 2017. 46(6): p. 957–967.

54. Moulson, A.J., et al., Diversity of Reactive Astrogliosis in CNS Pathology: Heterogeneity or Plasticity? Frontiers in Cellular Neuroscience, 2021. 15.

55. Linnerbauer, M. and V. Rothhammer, Protective Functions of Reactive Astrocytes Following Central Nervous System Insult. Frontiers in Immunology, 2020. 11.

56. Schiweck, J., B.J. Eickholt, and K. Murk, Important Shapeshifter: Mechanisms Allowing Astrocytes to Respond to the Changing Nervous System During Development, Injury and Disease. Frontiers in Cellular Neuroscience, 2018. 12.

57. Lian, H., et al., NFκB-activated astroglial release of complement C3 compromises neuronal morphology and function associated with Alzheimer’s disease. Neuron, 2015. 85(1): p. 101–115.

58. Chiareli, R.A., et al., The Role of Astrocytes in the Neurorepair Process. Front Cell Dev Biol, 2021. 9: p. 665795.

59. Duarte Azevedo, M., S. Sander, and L. Tenenbaum, GDNF, A Neuron-Derived Factor Upregulated in Glial Cells during Disease. J Clin Med, 2020. 9(2).

60. Celikkaya, H., et al., GATA3 Promotes the Neural Progenitor State but Not Neurogenesis in 3D Traumatic Injury Model of Primary Human Cortical Astrocytes. Front Cell Neurosci, 2019. 13: p. 23.

61. Kizil, C., et al., Regenerative Neurogenesis from Neural Progenitor Cells Requires Injury-Induced Expression of Gata3. Developmental Cell, 2012. 23(6): p. 1230–1237.

62. Requejo, C., et al., Morphological Changes in a Severe Model of Parkinson’s Disease and Its Suitability to Test the Therapeutic Effects of Microencapsulated Neurotrophic Factors. Molecular Neurobiology, 2017. 54(10): p. 7722–7735.

63. Katiyar, K.S., et al., Mechanical elongation of astrocyte processes to create living scaffolds for nervous system regeneration. J Tissue Eng Regen Med, 2017. 11(10): p. 2737–2751.

64. Pekna, M. and M. Pekny, The Complement System: A Powerful Modulator and Effector of Astrocyte Function in the Healthy and Diseased Central Nervous System. Cells, 2021. 10(7): p. 1812.

65. Kanmogne, M. and R.S. Klein, Neuroprotective versus Neuroinflammatory Roles of Complement: From Development to Disease. Trends Neurosci, 2021. 44(2): p. 97–109.

66. Morales, I., et al., Astrocytes and retrograde degeneration of nigrostriatal dopaminergic neurons in Parkinson’s disease: removing axonal debris. Translational Neurodegeneration, 2021. 10(1): p. 43.

67. Marcel, M., et al., Complement C3 Deficiency Leads to Accelerated Amyloid β Plaque Deposition and Neurodegeneration and Modulation of the Microglia/Macrophage Phenotype in Amyloid Precursor Protein Transgenic Mice. The Journal of Neuroscience, 2008. 28(25): p. 6333.

68. Wyss-Coray, T., et al., Prominent neurodegeneration and increased plaque formation in complement-inhibited Alzheimer’s mice. Proceedings of the National Academy of Sciences, 2002. 99(16): p. 10837–10842.

69. Rocha, E.M., B. De Miranda, and L.H. Sanders, Alpha-synuclein: Pathology, mitochondrial dysfunction and neuroinflammation in Parkinson’s disease. Neurobiology of Disease, 2018. 109: p. 249–257.

70. Emmanouilidou, E., et al., Assessment of α-Synuclein Secretion in Mouse and Human Brain Parenchyma. PLOS ONE, 2011. 6(7): p. e22225.

71. He-Jin, L., P. Smita, and L. Seung-Jae, Intravesicular Localization and Exocytosis of α-Synuclein and its Aggregates. The Journal of Neuroscience, 2005. 25(25): p. 6016.

72. Chen, K., et al., LRP1 is a neuronal receptor for α-synuclein uptake and spread. Molecular Neurodegeneration, 2022. 17(1): p. 57.

73. Morizawa, Y.M., et al., Reactive astrocytes function as phagocytes after brain ischemia via ABCA1-mediated pathway. Nature Communications, 2017. 8(1): p. 28.

74. Liddelow, S.A., et al., Neurotoxic reactive astrocytes are induced by activated microglia. Nature, 2017. 541(7638): p. 481–487.

75. Tsunemi, T., et al., Astrocytes Protect Human Dopaminergic Neurons from α-Synuclein Accumulation and Propagation. J Neurosci, 2020. 40(45): p. 8618–8628.

